# Thermodynamics and unbinding kinetics of A22 at multiple actin binding sites revealed by enhanced sampling simulations

**DOI:** 10.1101/2025.07.09.663866

**Authors:** Anuj Kumar, Debabrata Pramanik

## Abstract

The cytoskeletal protein plays a major role in various cellular processes. Understanding the interactions of small molecules with cytoskeletal protein therefore might help in the development of therapeutics. Employing combined molecular docking, all-atom molecular dynamics (MD) and infrequent well-tempered metadynamics (iWT-MetaD) we studied bacterial inhibitor A22 and actin protein interaction. Five probable A22 binding sites (S1-S5) were observed in actin and unbiased MD simulations of 100 ns – 500 ns showed structural stability at these sites. Interaction analyses showed A22 to mostly forms transient interactions except at sites S4 and S5 where long-lives interactions were present. Enhanced sampling simulations quantitatively estimated ligand dissociation free energy (Δ*G*) ∼ – 2 kcal/mol to – 3.5 kcal/mol for sites S1-S4 with residence times ∼ µs to ms. For site S5, we observed the highest binding affinity (Δ*G* ∼ – 6.05 ± 0.48 kcal/mol) and longest residence time ∼ 0.75 sec. Analyses of the dissociation trajectories predict multiple dissociation pathways for A22 and found key gatekeeper residues at specific-sites facilitating ligand dissociation. Comparison of A22 with other known inhibitors suggest that A22 binds actin with relatively lower affinity, however, exhibits site-specific unbinding kinetics. Thus, our study provides a detailed mechanistic overview of A22-actin interaction, its various binding modes, and unbinding kinetics. It also shows the importance and usage of infrequent metadynamics in exploring rare events like ligand-protein interaction. Deeper insights gained from this study expands our understanding of cytoskeletal ligand dynamics. These knowledges will be paramount in designing drug targeting cytoskeletal protein actin.

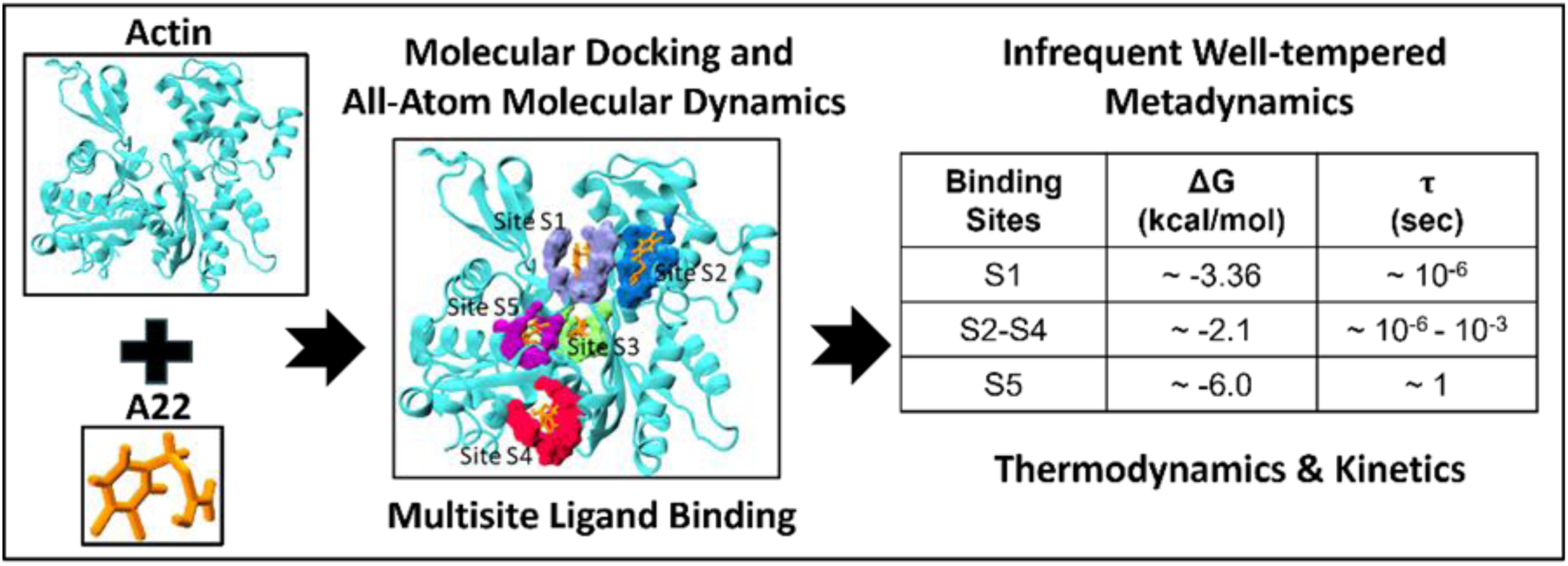
Graphical Abstract.

## Introduction

The cytoskeleton, in a biological cell, forms a complex network of protein filaments across the cytoplasm. These protein filaments are made up of three components: the microtubules, intermediate filaments, and microfilaments. Altogether, these components assist the cell in carrying out essential cellular functions.^1,2^ The actin protein, which belongs to the microfilament family, is extensively present and evolutionarily conserved in most eukaryotic cells. The three-dimensional (3D) structure of actin is globular in shape, with a sequence made up of 375 amino acid residues. It consists of four subdomains (SD): SD-1, SD-2, SD-3, and SD-4. A schematic of the actin crystal structure with four subdomains is shown in Figure 1. These subdomains in actin embed two large sites. There is a nucleotide-binding site between SD-2 and SD-4. The adenosine nucleotides, adenosine triphosphate/adenosine diphosphate (ATP/ADP), bind at this specific site together with a divalent positive ion (Mg^+2^ or Ca^+2^). The other site, known as the target binding site, located between SD-1 and SD-3, facilitates the binding of actin-binding proteins (ABPs) or small chemical compounds to this site. The actin protein can transform from a monomer (G-actin) to a polymer (F-actin) form. This transformation from one form to another helps actin in the execution of various cellular activities, like maintaining cell shape, cell polarity, cell division, cell migration, and intracellular transport processes.^3^ As actin is involved in diverse cellular functions, it is usually avoided as a target in drug therapeutics. Nevertheless, there are various studies that reported the involvement of the actin cytoskeleton in many pathological processes, such as tumorigenesis, invasion, and developmental diseases. Hence, small molecules that target actin and have the potential to inhibit actin functionalities hold promises in treating such kinds of diseases.^4–7^ Small molecules can bind to actin in either form, G-actin or F-actin, directly, and (de)stabilize the monomers or filaments, leading to inhibition of (de)polymerization.^8^ Previous studies reported that Phalloidin and Jasplakinolide bind to F-actin at the junction of three actin subunits, incorporate stability to F-actin, and thereby prevent depolymerization.^9^ Contrarily, Latrunculin A and Cytochalasin D bind between SD-2 and SD-4, close to the actin nucleotide-binding site^10^ and a hydrophobic site lying between SD-1 and SD-3, respectively. These molecules stabilize G-actin and prevent polymerization.^11,12^ Aplyronine A binds to a target-binding site (a hydrophobic cleft) between SD-1 and SD-3 of actin and inhibits the polymerization of G-actin to F-actin and depolymerizes F-actin to G-actin. Kabiramide C and Mycalolide B also bind to actin between SD-1 and SD-3 and act as actin-capping proteins to sever F-actin.^13,14^ Therefore, such small molecules are important in studying the actin cytoskeleton and can be used as therapeutics in various pathogenic conditions.

**Figure 1:**
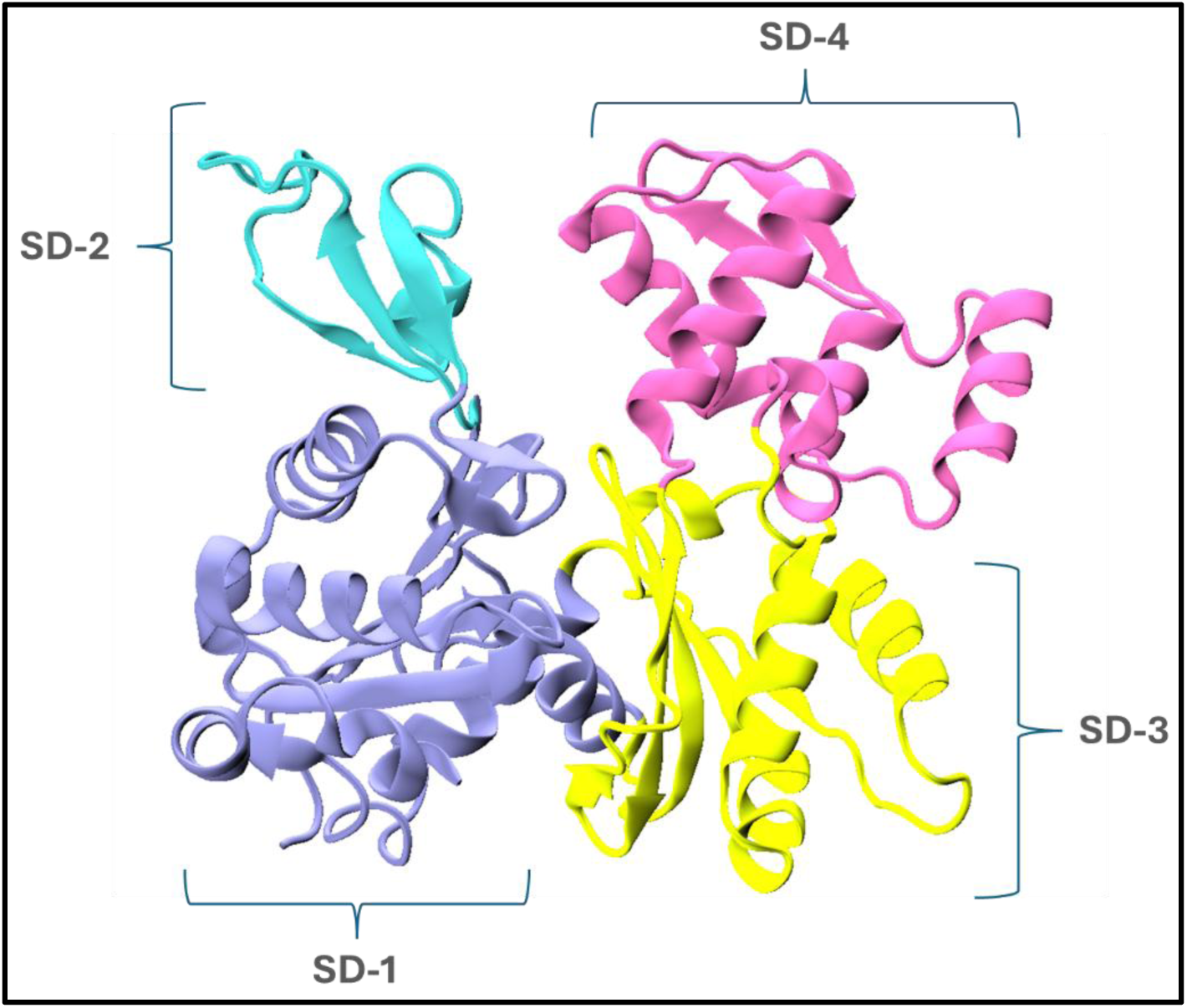
A schematic representation of the 3D structure of actin protein showing four subdomains, SD-1 (blue), SD-2 (cyan), SD-3 (yellow), and SD-4 (magenta).

The MreB protein, a bacterial homolog of eukaryotic actin, is present in almost all rod-shaped bacteria. Like actin, MreB also plays an important role in a broad range of cellular functions in bacterial cells. Although MreB lacks sequence similarity with actin, they are structural homologs.^15–19^ To date, many small-molecules have emerged as potential inhibitors (artificial/natural) of MreB, like benzyl-isothiourea derivatives (A22/MP265),^20,21^ small-molecule indole-class compounds (CBR-4830) and its derivatives, and TXH11106.^22,23^ Previous studies showed that A22 (S-(3,4-Dichlorobenzyl) isothiourea) does not exhibit significant cytotoxic effects on human peripheral blood mononuclear cells (PBMCs) and polymorphonuclear neutrophils (PMNs)^24,25^. In a recent study, Kumar et al. showed, through a combined computational and experimental study, that A22 interacts with eukaryotic actin by demonstrating its profound effects on actin organization in human glioblastoma cells^26^. It is important to note that although A22 is an inhibitor for MreB, it shows some finite amount of interaction with eukaryotic actin, though with lower affinity, and it raises the question of whether A22 can work as an inhibitor to actin or not. Therefore, a detailed understanding of the actin-A22 complex, its structural stability, preferential binding sites, key interactions facilitating the binding, quantitative estimation of binding affinities, and kinetic and mechanistic details would help to elucidate its role in actin inhibition. It is worthwhile to mention here that to date, the actin-A22 crystal structure has not been resolved. Therefore, the binding of actin with A22 and its binding mechanism have not been explored much. Hence, finding out the potential of A22 as an actin inhibitor requires attention^27^.

In the present study, we investigate the actin–A22 complex using all-atom molecular simulations and evaluate its structural and mechanistic details. Employing molecular docking and subsequent molecular dynamics simulations, we first explore various possible binding sites of A22 within actin. Docking results display that A22 can bind at multiple sites of actin. To get an idea of binding strength, we quantitatively estimated the dissociation free energy through enhanced sampling (well-tempered metadynamics) at each binding site. Using infrequent well-tempered metadynamics simulations, we calculate the ligand residence time for these binding sites. We elucidate the ligand dissociation pathways of A22 from the different binding sites of actin, with key residues along the dissociation paths facilitating the dissociation processes.

## Computational Methodologies

### Protein-ligand structure preparation

The initial crystal structure of the actin protein was obtained from the PDB database (PDB ID: 1YAG).^28^ Missing amino acid residues –-1MET, 2ASP, 3SER-– and missing atoms present in residues 40HIS, 41GLN, 43ILE, and 45VAL of the crystal structure were modelled using the CHARMM-GUI input generator tool.^29,30^ The initial coordinates of the A22 ligand was taken from the Cc-MreB (PDB ID: 4CZG) crystal structure.^31^ The structure of A22 was optimized using Gaussian software (Gaussian 16).^32^ To perform quantum optimization, a PM6 level calculation was followed by M06 level theory, using the 6-311G(d,p) basis set and a water dielectric medium as the continuum model. Single point RESP charges were calculated to obtain the partial atomic charges of A22.^33^

### Molecular docking strategy

We performed rigid receptor blind docking simulations using the AutoDock Vina 1.2.5 tool.^34,35^ To prepare the protein structure, hydrogens were first added to the protein. Subsequently, Gasteiger charges were added to actin. Similarly, for the ligand, we started with an optimized structure obtained from quantum optimization and assigned Gasteiger charges. Appropriate atom-types were assigned to both the protein and the ligand for molecular docking. A square grid box of size 126 along all three directions, with a spacing of 0.35 Å and centred at (9.172 Å, 6.280 Å, 14.098 Å), was prepared with energy range of 3 and an exhaustiveness of 32. We performed 100 independent docking runs for apo-actin, with each run consisting of 20 poses. Subsequently, only the top scoring docking poses were selected to compute the binding probability at each binding site. We used VMD and PyMOL to visualize and analyse the docking results ^36,37^.

### Molecular dynamics protocol

We carried out classical all-atom molecular dynamic simulation starting with the docked complex using the GROMACS-2022 software packages^38^. To model the interaction between the protein–ligand system, we used the all-atom CHARMM36 force field^39^ for the protein. To model A22-ligand interactions, we used the SwissParam web server to develop the force field parameters for A22.^40^ Water interactions were modelled using the TIP3P water model^41^. To neutralize the system, we added NaCl counterions. To compute bulk properties, periodic boundary conditions (PBCs) were applied in all three directions of the system. For system preparation, we first solvated the protein–ligand complex in an aqueous environment and neutralized the charge by adding counterions. To remove any overlap of water molecules with the solute, we performed energy minimization using the steepest descent algorithm. Subsequently, the system was simulated under NVT and then NPT ensembles to equilibrate it at room temperature (300 K) and atmospheric pressure (1 bar). We used the velocity rescaling thermostat and Parrinello-Rahman barostat to maintain the temperature^42^ and pressure^43^, respectively. To calculate electrostatic interactions, we employed the Particle-Mesh Ewald (PME) method. The Verlet cutoff scheme was used with a 1.4 nm cutoff for short-range electrostatic and van der Waals interactions.^44,45^ Using the LINCS algorithm, we constrained covalent bonds involving hydrogen atoms and heavy atoms, allowing the use of an integration time step of 2 fs with the leap-frog MD integrator. Once the system was equilibrated, we performed a production run to calculate various structural and interaction properties.

### Interaction fingerprint analysis

To explore the interactions between the ligand and protein in the bound state, we calculated the interaction fingerprints^46^ using ProLIF, a Python library^47^ from the unbiased MD simulation trajectories. ProLIF uses SMARTS patterns to describe molecular interactions between the ligand and the protein. In calculating the interaction fingerprints, two groups of atoms (one in the protein and the other in the ligand) were selected using SMARTS patterns. We calculated interactions such as hydrogen bonds, hydrophobic interactions, van der Waals, anionic, cationic, and cation-π interactions. All these interactions were determined based on different geometrical and atomic criteria. In the Supplementary Information, we provided the detailed criteria for various interactions, and the interaction fingerprint plots (shown in Figure S1(a-e)). We considered only those interactions between the ligand and the protein as stable if they persisted for more than 90% of the total simulation time.

### Infrequent well-tempered metadynamics

To calculate the free energy of dissociation of the ligand from the protein binding sites, we employed the metadynamics simulation technique, an advanced sampling method. Our interest was in both the dissociation free energy and its kinetics. Therefore, to extract the dissociation free energy from the metadynamics simulation, we used a variant of metadynamics known as Well-Tempered Metadynamics (WT-MetaD).^48–51^ In well – tempered metadynamics, a time dependent bias is deposited along a pre-selected reaction coordinate (RC), which helps the system escape from energy minima. By depositing the bias infrequently--i.e., by performing infrequent well-tempered metadynamics (iWT-MetaD) –-we can also extract kinetic details from a biased simulation^52–56^. The results of a WT-MetaD simulation depend strongly on the choice of the reaction coordinate (RC). A good RC should be able to demarcate the various metastable states in the system. Moreover, in iWT-MetaD, another crucial assumption is that the frequency of bias deposition must be such that the bias is not deposited in the transition state. This ensures that the transition state remains unperturbed while the system moves from one free energy well to another.

We used PLUMED,^57,58^ patched with GROMACS, to perform the WT-MetaD simulations. In our calculations, we used the distance between the centre of mass (COM) of the protein site residues and the ligand heavy atoms as the reaction coordinate (RC). In supplementary Figure S2 we provide the details of the protein residues in the binding sites and the ligand atoms selected for the RC in each case. To calculate distance (RC), we selected two groups, ligand heavy atoms centre of mass (COM) and COM of the protein residues in the binding site. To select protein COM, the following residues were chosen: site S1 (GLY15, PHE31, GLN59, TYR69, HSD73, GLY158, ASP184, ILE208, ARG210, LYS213), site S2 (ASP184, ASP187, LEU180, PRO264, SER271, LEU267), site S3 (ASN12, SER14, HSD73, GLU107, ASP157, THR160, SER300, VAL339), site S4 (VAL134, SER135, VAL 139, TYR143, SER170, PRO172, LEU346, PHE375), and site S5 (ASP11, ILE71, GLY75, VAL76, ASN78, TRP79, GLU83, TRP86, HIS87, THR106, ALA108, MET110, PRO112, ASN115, MET119, THR120, MET126), respectively. The parameters (hill height, width, deposition frequency, and bias factor) used in the metadynamics simulations were carefully optimized to ensure that the resulting statistics were independent of parameter choice. The details of the optimization process are provided in the supplementary information (Figure S3). The following parameters were used in the iWT-MetaD simulation: a gaussian bias (V_b_(*s*, *t*)) with a height of 0.8 kJ, a width of 0.01 nm, a bias factor of 12, and a temperature of 300 K. The bias was deposited every 8 ps (i.e. every 4000 steps) to maintain its infrequent nature. Utilizing reweighting technique^59,60^, we extracted free energy along the RC.

While computing average free energy, we first calculated individual probabilities, followed by the averaged probability, and then the average free energy using the following equations:

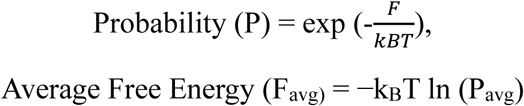

The error bar in free energy has been calculated by using the following equation.

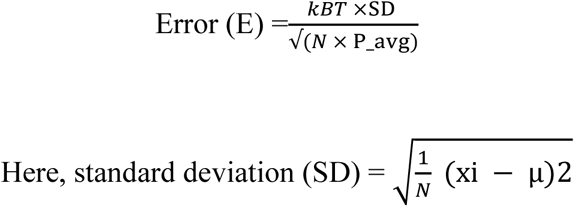

*N* is the number of samples, *x*_i_ is the random variable, and μ is the mean/average.

The individual free energy plots for all the five binding sites are provided in Figure S4.

### Residence time calculation

To calculate the transition time or ligand residence time (i.e., the amount of time a ligand takes to cross the transition state from the minimum of the energy landscape), we utilized Tiwary’s algorithm,^48^ which is based on the transition state theory. To obtain the unbiased residence time (τ_unbiased_), we first calculated the metadynamics-associated transition time (τ_biased_) and then computed the unbiased time (τ_unbiased_) using the following relation:

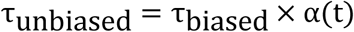

The acceleration factor (α) is calculated by the following equation.

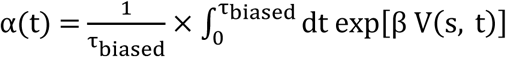

Where V(s, t) is the instantaneous bias potential deposited in the basin region along the reaction coordinate (RC). Here, β = 1/k_B_T, where k_B_ and T denote the Boltzmann constant and simulation temperature, respectively. To calculate the biased residence time (τ_biased_), we identified the individual transition times from the abrupt change in the sign of the first derivative of α with respect to simulation time. We calculated d(α(t))/dt as a function of time t for each run, as shown in supplementary Figure S5. From these plots, the biased transition time (τ_biased_) was extracted by identifying the point of abrupt sign change in the derivative of α. We also plotted the time evolution of the collective variable as A22 dissociated from the binding site, as shown in Figure S6. To assess the accuracy of the calculated residence times from the infrequent metadynamics trajectories, we performed a Kolmogorov–Smirnov (KS) test and calculated the p-value for each system, as listed in Table 4. Theoretical details and plots related to the KS test are provided in supplementary Figure S7.

## Results and Discussion

### Detection of multiple ligand binding sites

As shown in Figure 1, the actin protein consists of four subdomains: SD-1, SD-2, SD-3, and SD-4. Based on its 3D structure, the actin protein can have multiple binding sites, either embedded within the protein or located on its surface. Since actin contains several ligand-binding regions, small chemical molecules can bind to these sites and influence the structural and functional properties of the protein. It is worth mentioning that previous studies have identified two predominant interdomain regions in actin: the nucleotide-binding site (between SD-2 and SD-4) and the target-binding site (between SD-1 and SD-3). To explore the potential binding sites in actin for a small chemical molecule like A22, we first performed molecular docking simulations to identify regions where A22 could bind. We carried out 100 independent docking simulations, with each run generating 20 different ligand poses. Figure 2(a) displays all 2000 docked poses collectively. As shown in the figure, A22 can bind to various sites on actin. Based on the docking results, we analysed the distribution of A22 across these sites. Figure 2(b) highlights all possible binding sites with the corresponding percentage occupancy of A22. Based on occupancy and docking scores (as shown in Figure 2(c)), we selected four binding sites, named S1, S2, S3, and S4. Table 1 provides the percentage of A22 binding to each site on the actin protein. To identify the most relevant binding sites, we included only those with an occurrence percentage greater than 10%, ensuring that only the more abundant sites were considered. To determine the protein residues forming each binding site, we defined a region within a 5 Å cutoff surrounding the ligand at each binding site and selected the protein residues present within this region (listed in Table S1). Site S1 is located between the SD-2 and SD-4 subdomains. Previous studies have reported that small molecule inhibitors such as Latrunculin-A/B bind to this site. Site S2 is located on the protein surface at the SD-4 subdomain. Site S3 lies between all four subdomains (SD-1, SD-2, SD-3 and SD-4). Site S4 is an important binding site located between SD-1 and SD-3 and is known to accommodate various ligands such as Cytochalasin D^11^, Kabiramide C^14^, Mycalolide B^61^, Reidispongiolide A^62^, Sphinxolide^62^, Aplyronine A^63^, and Chivosazole A^64^. It should be noted that sites S2 and S5 have not been previously reported as ligand-binding sites, to the best of our knowledge, and can be considered as new binding sites in the actin protein. Molecular docking analysis indicates that A22 exhibits significant binding at all these sites, as shown in Table S1.

**Figure 2:**
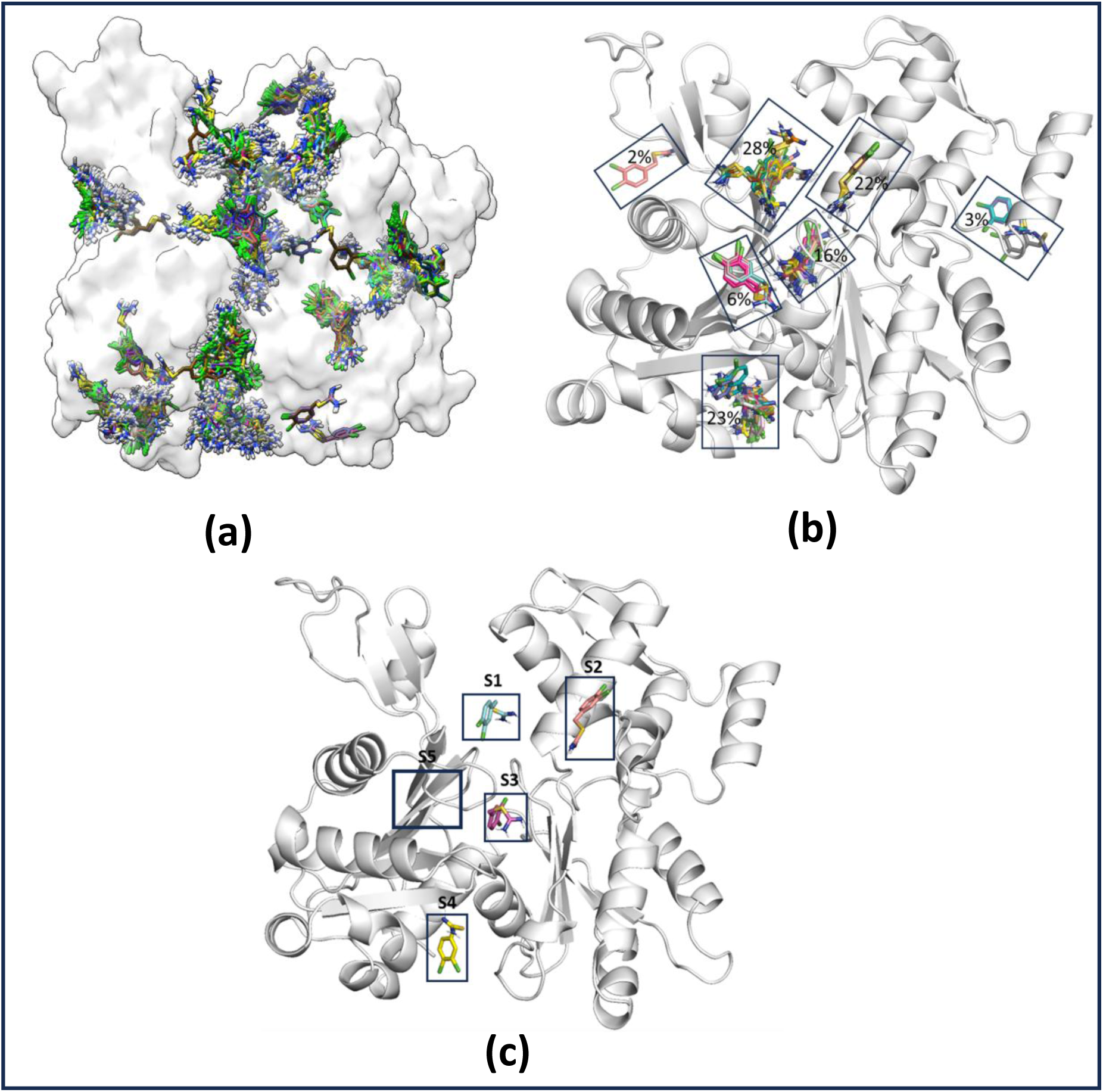
Instantaneous snapshots of the A22-actin binding poses. (a) 2000 molecular docked poses of the binding of A22 with actin protein at different binding sites. These binding poses are from 100 independent molecular docking simulations, each individual run consisting of 20 ligand poses. (b) We showed few selected binding sites based on the percentage of A22 binding. (c) Actin in apo form has 5 binding sites named as S1, S2, S3, S4, and S5. The respective percentage of a22 binding at these sites are as follows: site S1 28%; site S2 22%; site S3 16%; site S4 23%. The remaining sites having a22 binding percentage of less than 10% have not been included here.

**Table 1:** The table shows docked systems studied, number of independent docking runs, number of binding sites > 10 % occurrence, name of the sites and their percentage of occurrence.

| Protein | Ligand | Total number of independent docking runs | Number of binding sites with > 10% | Name of binding sites | Binding percentage (%) |
| --- | --- | --- | --- | --- | --- |
| Actin | A22 | 100 | 4 | S1 | 28 |
|  |  |  |  | S2 | 22 |
|  |  |  |  | S3 | 16 |
|  |  |  |  | S4 | 23 |

Previous studies have shown that actin and its bacterial homolog, MreB, share structural homology. The MreB crystal structure reveals that the bacterial inhibitor A22 binds to a site which is distinct from the four aforementioned ones--adjacent to the nucleotide-binding site^31^(Figure 2c)). Therefore, we included this binding site as an additional potential site for A22 in actin, naming it as site S5. Since site S5 is buried deep within the protein’s 3D structure, blind rigid molecular docking simulations could not assess this site. To obtain an initial bound pose at this location, we manually placed A22 by translating and rotating it to align with the conformation observed in MreB. Figure 2(c) illustrates all five binding sites within the four subdomains of actin. Since molecular docking does not account for the contributions of the aqueous environment, counterions, or system dynamics, we further conducted all-atom molecular dynamics (MD) simulations for each of the five binding sites. Through unbiased MD simulations, we calculated various structural properties to assess the stability of the actin-A22 complex at each of these binding sites.

### Structural stability analysis

In Table 2, we provide details of the unbiased MD simulations and associated system information. We performed 100 ns to 500 ns of unbiased MD simulations for each of the five systems. For each system, two independent unbiased MD simulations were carried out and the resulting trajectories were analysed. To assess the structural stability of the protein-ligand complexes with A22 bound at specific binding sites of the actin protein, we calculated the root mean square deviation (RMSD), root mean square fluctuation (RMSF), and radius of gyration (Rg) of the protein backbone atoms (C, C_α_, N, and O) from the unbiased MD trajectories. These analyses help to identify the regions of high fluctuation, assess conformational stability, and evaluate the compactness of the protein in the presence of the A22 ligand at the different binding sites: S1, S2, S3, S4, and S5. In Figure 3, we show the RMSD and Rg as a function of time, and RMSF as a function of residue number. As observed in Figure 3(a-e), the RMSD of protein remains around ∼ 0.2 – 0.25 nm over time, with fluctuations within ∼ 0.1 nm across all systems. The root means square fluctuation (RMSF) of protein residues, when bound to A22, reflects the local fluctuations of residues during the simulations. No significant conformational changes were observed while binding to actin sites except at S5. At site S5, we noticed a slight reduction in the RMSF, specifically in the DNase-I binding loop (D-loop) region. This region corresponds to the SD-2 subdomain of actin. To further evaluate the compactness of the complexes during the simulations, we computed the radius of gyration (R_g_) over time (Figure 3(a-e)). The R_g_ values ∼ 2.25 nm remains constant throughout the simulations, indicating that the structures remained compact and stable. overall, the actin-A22 complexes were well equilibrated, allowing us to proceed with further analysis to identify key interactions, crucial residues, and other important features of the A22-actin binding mechanism.

**Figure 3:**
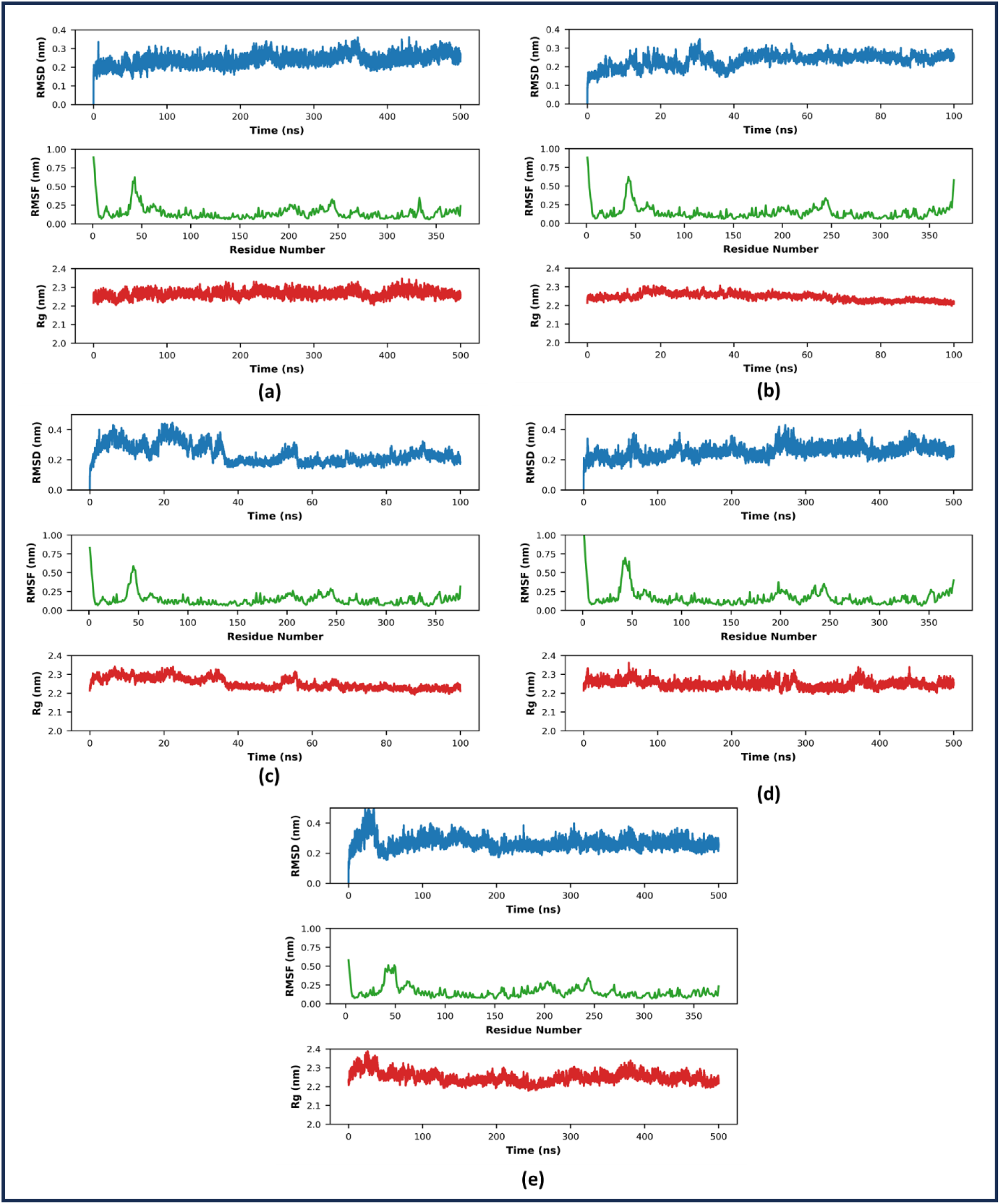
Time evolution of various structural properties for A22-actin system for sites S1, S2, S3, S4 and S5. We calculated RMSD (root mean square deviations), RMSF (root mean square fluctuations), and Rg (radius of gyration) with time. The RMSD, RMSF and Rg are shown for binding site S1 in (a), binding site S2 in (b), binding site S3 in (c), binding site S4 in (d), and binding site S5 in (e). For site S1, S4 and S5 we performed 500 ns of unbiased MD simulations, whereas for site S2 and S3, 100 ns of unbiased simulations have been performed.

**Table 2:** The table shows details of unbiased simulations: different systems studied, box sizes, total atoms in the systems, number of water molecules, total counterions (Na+ and Cl-), and duration of unbiased MD simulations.

| System | Site | Box sizes<br>(X nm, Y nm, Z nm) | Total number of atoms in the system | Number of water (H<sub>2</sub>O) molecules | Number of sodium (Na<sup>+</sup>) and chlorine (Cl<sup>-</sup>) ions | Duration of unbiased simulations (ns) |
| --- | --- | --- | --- | --- | --- | --- |
| Actin + A22 | S1 | 9.62, 9.62, 9.62 | 91327 | 28454 | 67, 56 | 500 |
|  | S2 | 9.62, 9.62, 9.62 | 91306 | 28447 | 67, 56 | 100 |
|  | S3 | 9.62, 9.62, 9.62 | 91318 | 28451 | 67, 56 | 100 |
|  | S4 | 9.62, 9.62, 9.62 | 91321 | 28452 | 67, 56 | 500 |
|  | S5 | 9.49, 9.49, 9.49 | 87622 | 27231 | 63, 53 | 500 |

### Interactions facilitating A22 – actin binding

To identify the various types of interactions and key interacting residues involved, we carried out interaction fingerprint analyses for all systems based on the unbiased MD simulation data. In supplementary information, Figure S2(a-e) presents the interaction fingerprint plots for each system. Based on the duration of these interactions, we classified them into two categories: stable/long-lived, interactions surviving for at least 90% of the simulation time; and transient/short-lived, interactions occurring for at least 30% but less than 90%. In Figure 4(a-e), we present the 2D interaction maps for interactions that persisted for more than 30% of the simulation time across all five binding sites. A list of residues involved in the actin-A22 interactions at each binding site is provided in Table 3. At site S1, A22 forms a stable hydrophobic interaction with residue TYR69. Once this interaction breaks, the ligand A22 quickly dissociates from the site. For site S2 (lying on the surface of the protein) and site S3 (positioned between all four subdomains of actin), we did not observe any stable (long-lived) interactions contributing to binding. At site S4, A22 forms stable hydrophobic and VdW interactions with LEU346. Even after 500 ns of unbiased MD simulation, the ligand remains bound at this site. Similarly, at site S5, A22 establishes several stable hydrophobic and VdW interactions with ALA108, VAL76, ILE71, ASN12, consistent with findings previously reported by Kumar et al^26^. Since sites S2 and S3 lack long-lived interactions, it is possible that transient interactions are responsible for retaining the ligand at the binding sites during the simulation. For each site, we performed two independent unbiased MD simulations and analysed both trajectories to ensure robustness in our results. It is important to note that for sites S1, S2 and S3, A22 dissociates from the binding sites relatively quickly, within ∼ 100 ns or less. Therefore, only those snapshots where the ligand remains bound within the site were considered for computing the interaction fingerprints. A comprehensive list of all interacting residues and the nature of transient interactions is provided in Table 3.

**Figure 4:**
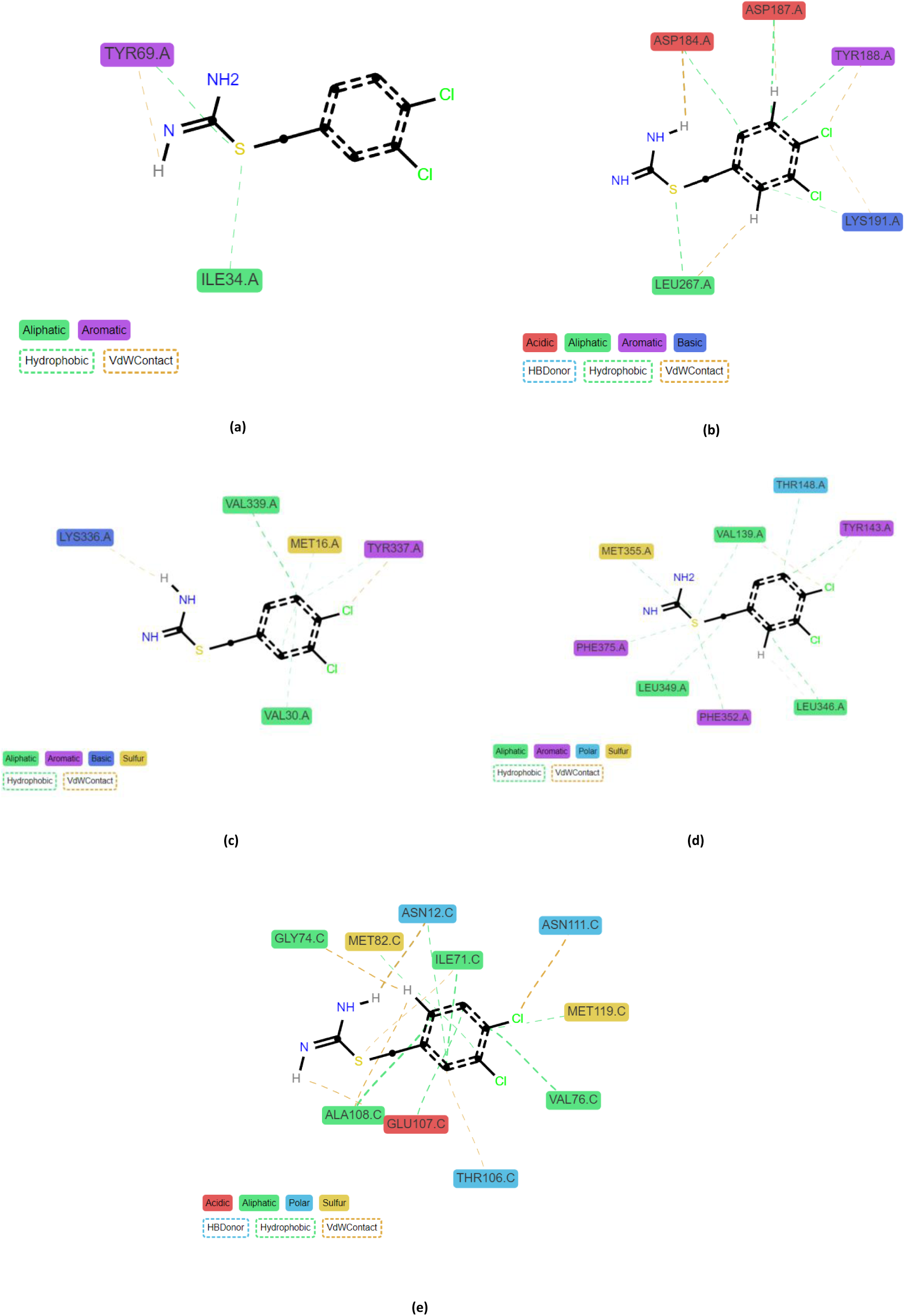
2D interactions plot of A22 with actin protein. (a) The 2D interaction maps for site S1 in actin, (b) site S2, (c) site S3, (d) site S4, and (e) site S5, respectively. We provided the types of interactions, like HB Donor, hydrophobic, VdW contacts by different colours also the protein residues forming these interactions in the 2D map. To calculate this 2D interaction map we used a cutoff of xxx.

**Table 3:**
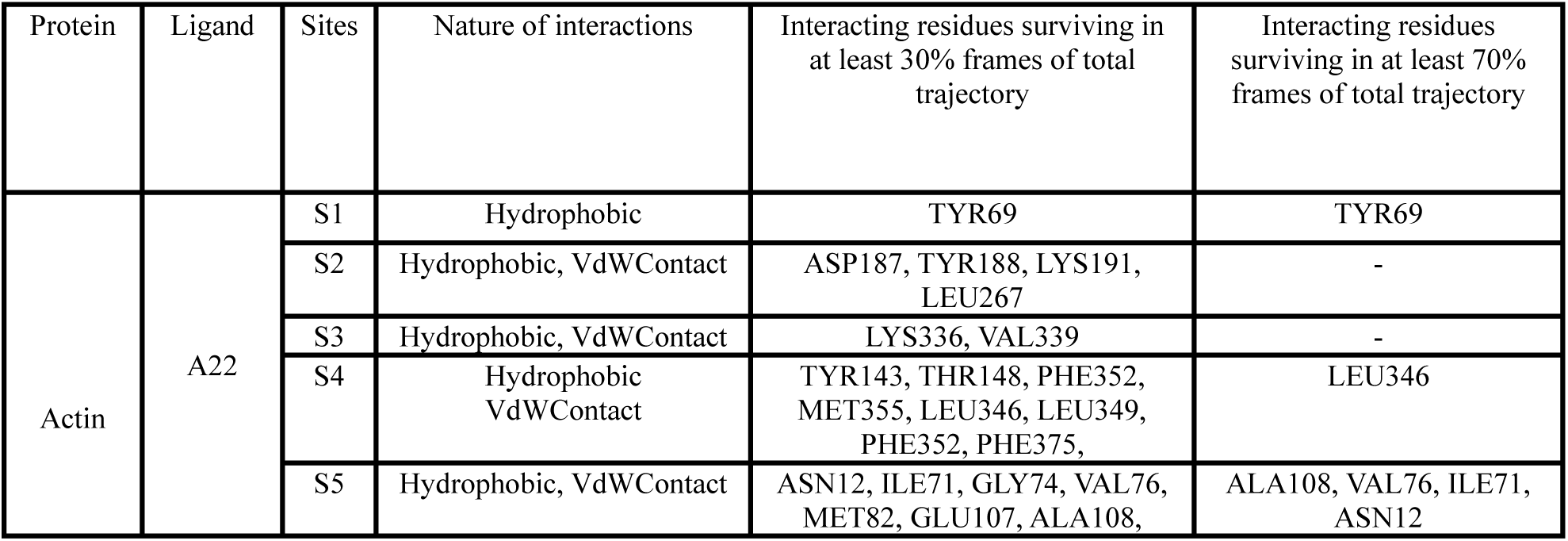
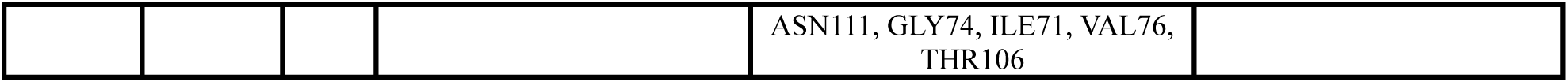
The table shows the type of interactions between A22 ligand and actin protein, along with a list of key residues, transient as well as stable protein residues, involved in the binding process at each of these five binding sites S1, S2, S3, S4, and S5, from molecular dynamics trajectories.

| Protein | Ligand | Sites | Nature of interactions | Interacting residues surviving in at least 30% frames of total trajectory | Interacting residues surviving in at least 70% frames of total trajectory |
| --- | --- | --- | --- | --- | --- |
| Actin | A22 | S1 | Hydrophobic | TYR69 | TYR69 |
|  |  | S2 | Hydrophobic, VdWContact | ASP187, TYR188, LYS191, LEU267 | - |
|  |  | S3 | Hydrophobic, VdWContact | LYS336, VAL339 | - |
|  |  | S4 | Hydrophobic VdWContact | TYR143, THR148, PHE352, MET355, LEU346, LEU349, PHE352, PHE375, | LEU346 |
|  |  | S5 | Hydrophobic, VdWContact | ASN12, ILE71, GLY74, VAL76, MET82, GLU107, ALA108, | ALA108, VAL76, ILE71, ASN12 |
|  |  |  |  | ASN111, GLY74, ILE71, VAL76,<br>THR106 |  |

**Table 4:** Number of independent infrequent metadynamics runs, average residence times, average characteristics times, p-values, and average dissociation free energies at sites S1, S2, S3, S4, and S5 respectively.

| Binding Sites | Number of Inf-MetaD Runs | Average Dissociation Free Energy (Kcal/mol) | Average Residence Time (s) | p-value |
| --- | --- | --- | --- | --- |
| S1 | 30 | - 3.36 ± 0.30 | $2.79 \times 10^{-6} \pm 0.005$ | 0.78 |
| S2 | 30 | - 2.10 ± 0.53 | $3.16 \times 10^{-5} \pm 9.78 \times 10^{-5}$ | 0.77 |
| S3 | 30 | - 2.09 ± 0.08 | $3.56 \times 10^{-3} \pm 0.010808$ | 0.75 |
| S4 | 30 | - 2.17 ± 0.37 | $1.84 \times 10^{-3} \pm 0.004744$ | 0.27 |
| S5 | 27 | - 6.05 ± 0.48 | $7.47 \times 10^{-1} \pm 2.281557$ | 0.27 |

After identifying the binding sites and the key interactions stabilizing A22 at each site on actin, we next aimed to quantify the binding affinity of A22 to actin at these respective sites. To this end, we first examined the dissociation behaviour of A22 through unbiased MD simulations.

### Unbiased dissociation events

We performed unbiased MD simulations over a timescale ranging from 100 ns to 500 ns. Structural properties like RMSD, RMSF, and R_g_ are useful in monitoring the overall structural changes in complex biomolecular systems. To explore the binding affinity of A22 at different actin binding sites, we observed the dissociation of A22 from these sites. To qualitatively observe the dissociation behaviour and quantitatively assess the dissociation free energy, we calculated centre-of-mass (COM) distance between all atoms of the ligand and the backbone atoms of the surrounding binding site residues over time. In Figure 5, we present the time evolution of the COM distance for all five binding sites. At site S1, A22 dissociates from the binding site at approximately130 ns, re-enters the same site, remains bound for ∼ 350 ns and then dissociates again. At site S2, a surface-exposed site, the ligand binds weakly and dissociates within approximately 50 – 100 ns. Similarly, at site S3, A22 exits the binding pocket within 100 ns of simulation time. These dissociation events suggest that A22 exhibits weaker binding affinity at sites S1, S2, and S3. In contrast, at site S4 and S5, we did not observe any dissociation events for A22, even after 500 ns of simulation. This indicates that A22 binds with comparatively higher affinity to sites S4 and S5.

**Figure 5:**
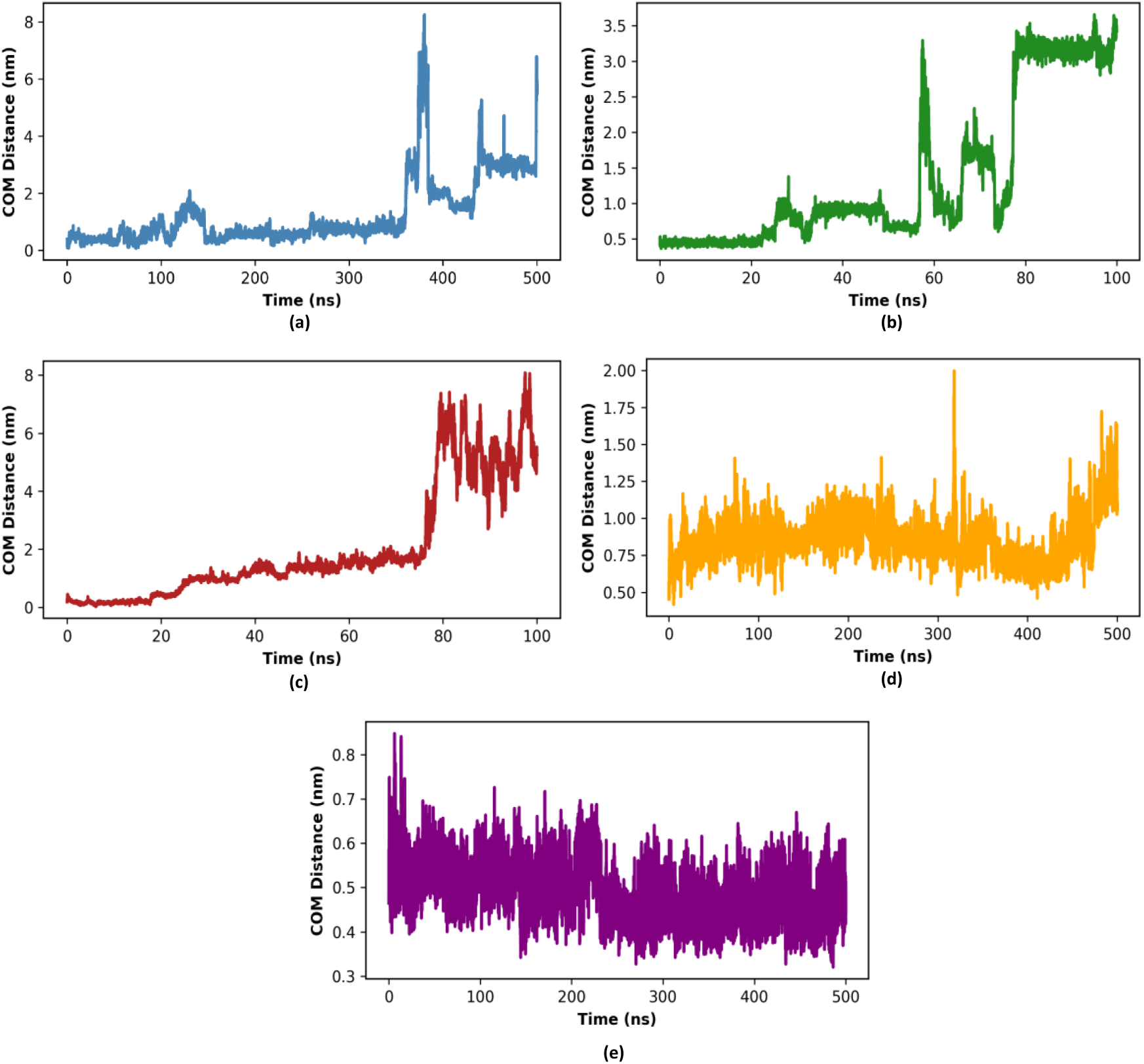
The plots show time evolution of the centre of mass distances for five binding sites. These are calculated from unbiased MD simulation trajectories. The binding sites are respectively (a) site S1, (b) site S2, (c) site S3, (d) site S4, and (e) site S5.

The dissociation of a ligand from a protein binding pocket is a rare event in nature. As observed from Figure 5, 100 ns to 500 ns of unbiased MD simulations were sufficient to capture ligand dissociation from sites S1, S2, and S3, whereas for sites S4 and S5, longer simulations beyond 500 ns would be required. To quantitatively assess the binding affinity of A22 at these specific binding sites, we therefore proceeded with enhanced sampling simulations at each site. Using enhanced sampling technique metadynamics, we quantitatively calculated the ligand dissociation free energy, dissociation pathways, and residence times.

### Ligand dissociation free energies

To quantitatively estimate the ligand dissociation free energy, we performed infrequent well-tempered metadynamics (iWT-MetaD) simulations for all five binding sites: S1, S2, S3, S4, and S5. In these simulations, the system was biased along a reaction coordinate (RC) defined as the distance between the centres of mass (COM) of the ligand and the selected binding pocket residues. Instantaneous snapshots showing the selected protein residues at the five binding sites and the ligand COM groups used as RC are schematically illustrated in supplementary Figure S3(a-e). The corresponding figure caption in the supplementary information lists the specific binding pocket residues chosen for the biased simulations. For each system, we performed 30 independent iWT-MetaD simulations and computed independent free energy profiles, shown in supplementary Figure S4(a-e). In Figure 6, we present the averaged dissociation free energy profiles as a function of the reaction coordinate, including error bars. The dissociation free energy (Δ*G*) was calculated as the difference between the global minimum and the transition state along the free energy profile. As shown in Figure 6, the Δ*G* values for A22 dissociation at each site are as follows: site S1 – 3.36 ± 0.30 kcal/mol; site S2 ∼ – 2.10 ± 0.53 kcal/mol; site S3 ∼ – 2.09 ± 0.08 kcal/mol; site S4 ∼ – 2.17 ± 0.37 kcal/mol, and site S5 ∼ – 6.05 ± 0.48 kcal/mol.

**Figure 6:**
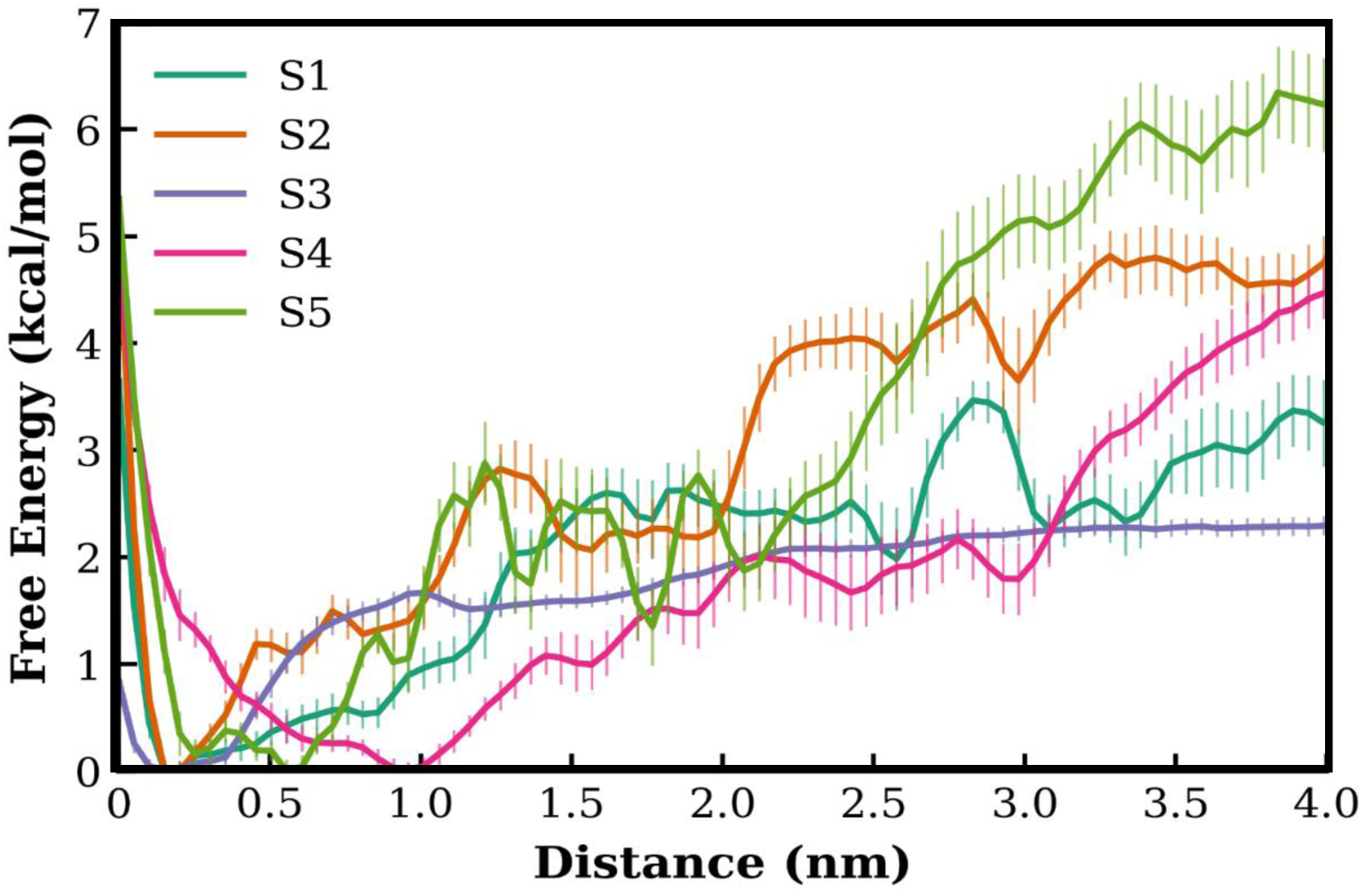
This shows the Free Energy Profile of actin+A22 system when ligand A22 dissociates from the binding sites S1, S2, S3, S4, S5. The error shows the standard deviation from the mean.

To assess the convergence of the free energy profiles, we calculated block averages for each case. Figure 7 shows the average error in the free energy profiles as a function of block size for all five systems. The results indicate good convergence, with the maximum error in the calculated free energies remaining below ∼ 0.5 kcal/mol. Details of the number of iWT-MetaD simulations performed and the averaged dissociation free energy values with associated errors for all five binding sites are provided in Table 4.

**Figure 7:**
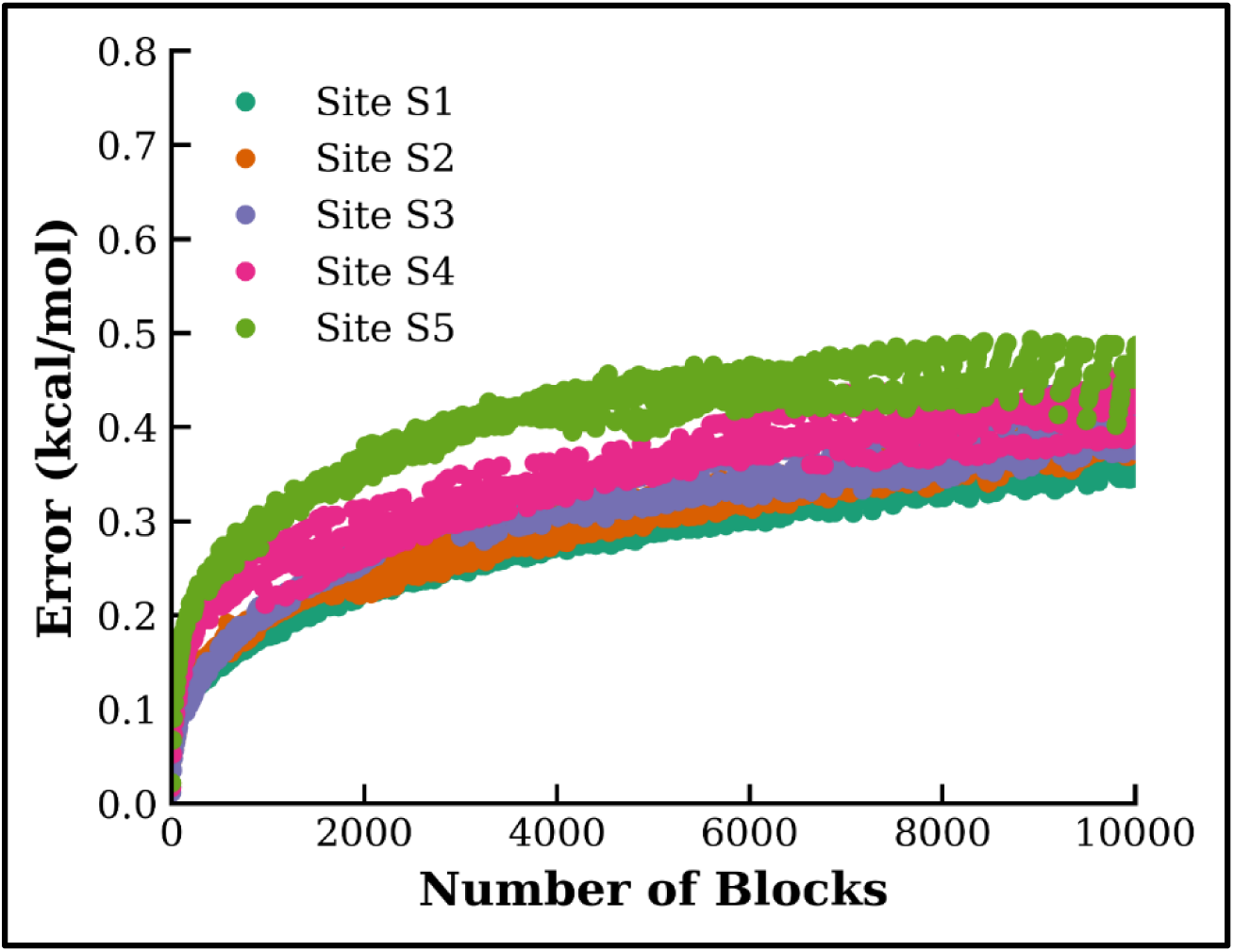
To check the convergence in the free energy calculation, we performed block analysis. In this plot, we showed average error in free energy profiles as a function of block size for all five systems. This shows the maximum error in free energies always remains less than ∼0.5 kcal/mol.

### Ligand residence times

The residence time of a ligand at any binding site refers to the duration the ligand remains within the binding site. In our study, we calculated the residence time as the average time a ligand takes to move from its energy minimum to the transition state region along the free energy surface. To calculate the biased dissociation time, we extracted the dissociation time from the time derivative of alpha (acceleration factor) vs time plots (as shown in Figure S5). From the RC vs time plots (Figure S6), we estimated the approximate time of ligand dissociation from the binding site by identifying when it crosses the transition state. For each of the five sites, we first calculated the individual residence time and then computed the averaged residence time, which is provided here. We performed the Kolmogorov-Smirnoff (KS) test to verify the reliability of the residence times calculated from our infrequent well-tempered metadynamics simulations. The test ensures that the deposited bias does not corrupt the transition state barrier, confirming that the calculated residence times are not affected by the deposited bias. Figure S7 presents the KS test plots for all five binding sites. Table 4 lists the averaged residence times and p-values. A p-value greater than 0.05 indicates that the deposited bias has not perturbed the transition state barrier.

For site S1, we estimated an average residence time of ∼ 2.79 x 10-6 ± 0.005 s, p-value 0.78; for site S2, average residence time ∼ 3.16 x 10-5 ± 9.78 x 10-5 s, p-value 0.77; for site S3, average residence time ∼ 3.56 x 10-3 ± 0.010808 s, p-value 0.75; for site S4, average residence time ∼ 1.84 x 10-3 ± 0.004744 s, p-value 0.27; for site S5, average residence time ∼ 7.47 x 10-1 ± 2.281557 s, p-value 0.27.

### Ligand dissociation pathways

Exploring ligand dissociation pathways is important in the development of novel drugs. Using trajectories from infrequent well-tempered metadynamics simulations, we analysed the independent pathways chosen by the ligand A22 while dissociating from each binding sites. In Figure 8, we show snapshots of ligand dissociation pathways for all five systems and provide the key protein residues along each path that facilitate ligand dissociation. A22 dissociates through two independent pathways from site S1, along one pathway from sites S2, S3, S4, and three independent pathways from site S5. Our further analysis elucidates the important protein residues interacting with ligand A22 while dissociating along respective independent paths from the binding sites. To calculate these residues, we used the following procedure. Along the dissociation path, once the ligand is 1 nm away from the centre of the binding sites, we start counting the interaction within 0.35 nm cutoff between ligand and protein residues considering ligand centre of mass as the origin. Then out of all these counted interactions, we considered only those interactions which exist for more than 50 % of the time. This time we considered from the point when ligand crossed initial 1 nm of distance along the specific path. Similar procedure we followed to calculate interactions for all other independent trajectories which dissociate along the same path. Later, from this collection of residues we selected only the common residues which were present in all these independent trajectories along any specific path.

**Figure 8:**
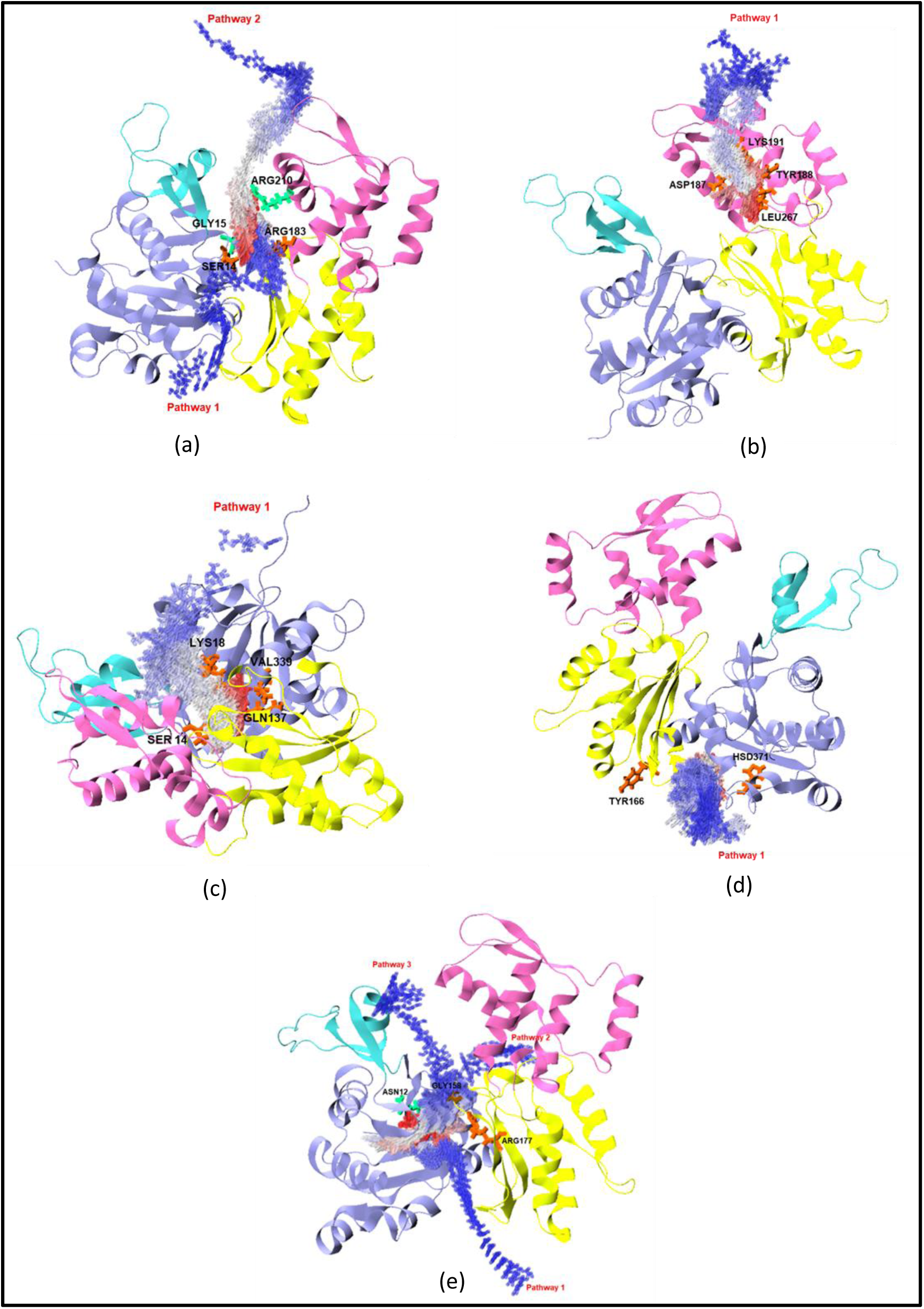
The figure shows schematic representation of the dissociation pathways for A22 ligand for five different binding sites in the actin protein. (a) For site S1, the ligand A22 can dissociate along two independent pathways as represented in the figure. (b) For site S2, we observed a single ligand dissociation pathway, (c) site S3, a single ligand dissociation pathway, (d) site S4, also we observed a single ligand dissociation pathway. (e) For site S5, we observed three independent dissociation pathways. Along each of these pathways, we also analysed the important protein residues which are helping in the dissociation process along the pathways and provided in the representation.

The following residues form major interactions and facilitate the dissociation processes.

For site S1: along pathway 1 – ARG183, SER14; pathway 2 – GLY15, ARG210.

For site S2: along pathway 1 – LEU267, TYR188, LYS191, ASP187.

For site S3: along pathway 1 – SER14, VAL339, GLN137, LYS18.

For site S4: along pathway 1 – TYR166, HSD371.

For site S5: along pathway 1 – ARG177; pathway 2 – ASN12; pathway 3 – GLY158. In Table 5, we provided the number of independent pathways and key residues along each of these paths for all five binding sites.

**Table 5:** The table shows the number of dissociation pathways for each of these binding sites, and the residues involved in the ligand dissociation along a particular dissociation pathway.

| Binding Sites | Number of dissociation pathways | Residues guiding the dissociation |
| --- | --- | --- |
| S1 | 2 | Pathway 1: ARG183, SER14; Pathway 2: GLY15, ARG210 |
| S2 | 1 | Pathway 1: LEU267, TYR188, LYS191, ASP187 |
| S3 | 1 | Pathway 1: SER14, VAL339, GLN137, LYS18 |
| S4 | 1 | Pathway 1: TYR166, HSD371 |
| S5 | 3 | Pathway 1: ARG177; Pathway 2: ASN12; Pathway 3: GLY158 |

### Corelation between free energy and residence time

After obtaining independent free energies and residence times of A22 for five binding sites (S1-S5), we explored how these two quantities correlate. The dissociation free energy and residence time of a ligand-protein system are related by the following basic equation that relates free energy (ΔG) and dissociation constant (K_d_):

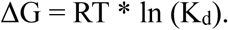

Here, K_d_ is defined as the ratio of dissociation rate constant over association rate constant:

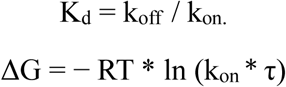

Where residence time, τ is given by

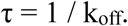

Since, τ and k_off_ are inversely related, therefore stronger binding affinity corresponds to a longer residence time, and vice-versa.

Figure 9 shows Δ*G* versus τ plot. For site S1, Δ*G* ∼ – 3.4 kcal/mol, τ ∼ 2.8 µs.

**Figure 9:**
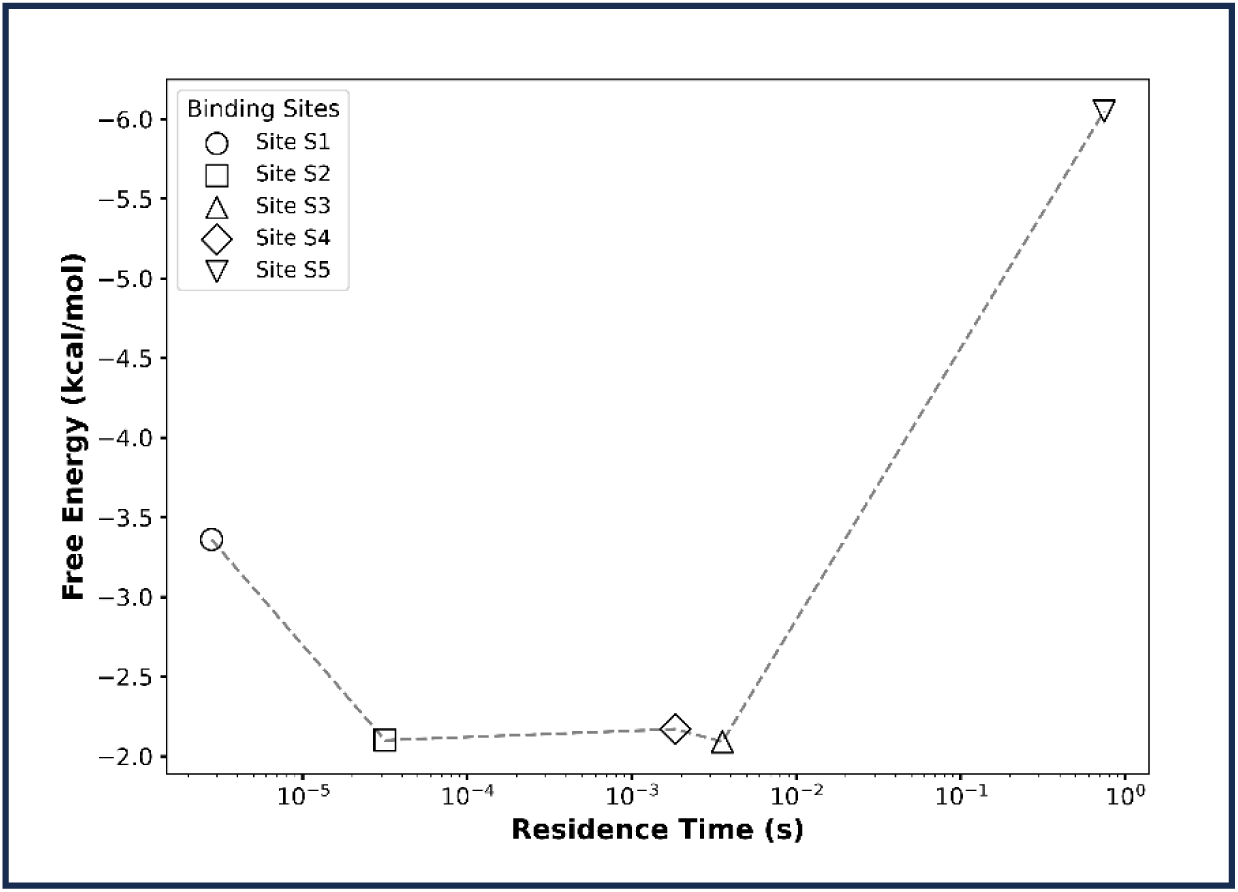
The plot shows correlation between “Free Energy” and “Residence Time” for actin and A22 system for binding sites S1, S2, S3, S4, and S5.

For site S2-S4, Δ*G* ∼ – 2.1 to –2.2 kcal/mol, and τ ∼ 30 µs, 3.5 ms, 1.8 ms, respectively, whereas for site S5, Δ*G* ∼ – 6.0 kcal/mol, and τ ∼ 0.75 s. Hence, Δ*G* ranges from ∼ –2 kcal/mol to –6 kcal/mol and τ ∼ µs to sec. Although Δ*G* and τ shows a good correlation, however, some deviation observed which can be attributed to enthalpic or entropic contributions influencing kinetics. Site S5 shows the highest binding affinity and longest residence rime which is consistent with the stronger ligand-protein interaction.

### Comparison with previously reported actin binding sites

As per reported studies on actin – inhibitors interactions, predominantly two major binding sites are predicted: the nucleotide-binding site and the target-binding site. From combined molecular docking and MD simulations, we additionally found out A22 to bind at some other regions on the protein as well. Here, we compare our observation with previously reported sites to validate our study.

Site S1: The binding site for Latrunculin A corresponds to this site. TYR69 is a key residue in both A22 and Latrunculin A binding. Latrunculin A has a dissociation constant of 0.2 µM (∼ – 9.19 kcal/mol). In comparison, A22 has a free energy of –3.36±0.30 kcal/mol at this site.

Site S4: The actin inhibitors such as MiuA, Reidispongiolide A, Cytochalasin D, Kabiramide C bind to site S4, the target-binding site. We observed common residues, including PHE352 (A22 and MiuA) and TYR143 (A22, MiuA, Kabiramide C). Kabiramide C has a K_d_ of 100 nM or less (–9.60 kcal/mol), and MiuA has a reported free energy of ∼ −39.4 kcal/mol. In comparison, A22 shows weaker free energy of – 2.17 ± 0.37 kcal/mol.

Details of binding sites, interacting residues, experimental and computational binding affinities are listed in Table 6.

**Table 6:** It shows reported results for actin and various ligand systems, number of binding sites, interacting restudies, experimental and computational binding affinities.

| Protein + Ligand Systems | Number of Binding Sites | Interacting Residues | Dissociation Constant (Experimental) | Free Energy (Computational) (kcal/mol) |
| --- | --- | --- | --- | --- |
| actin + A22 | S1 | TYR69 | - | - 3.36 ± 0.30 |
|  | S2 | - |  | - 2.10 ± 0.53 |
|  | S3 | - |  | - 2.09 ± 0.08 |
|  | S4 | LEU346 |  | - 2.17 ± 0.37 |
|  | S5 | ALA108, VAL76, ILE71, ASN12 |  | - 6.05 ± 0.48 |
| actin + MiuA<sup>65</sup> | M1 (This reported site corresponds to site S1) (as per our nomenclature) | Leu110, Ile136, Ile175, Val139, and Ala170 of actin, as well as Tyr133, Phe352, Tyr143 | - | -39.4 |
|  | M2 | Lys336 |  | -35.3 |
|  | M3 | Leu349 and Ile345 |  | -19.1 |
|  | M4 | Phe200 and Leu242 |  | -21.9 |
|  | M5 | Asp157 |  | -29.3 |
| actin+ Latrunculin A<sup>66</sup> | The reported site corresponds to site S1 | Gly15, Leu16, Pro32, Ile34, Gln59, Tyr69, Asp157, Gly182, | 0.2 μM | -9.19 |
|  |  | Arg183, Thr186,<br>Arg206, Glu207, and<br>Arg210 |  |  |
| actin+Reidispon<br>giolide A<sup>62</sup> | The reported site<br>corresponds to site<br>S4 | Ser141, ILE330, Pro332,<br>Ser338 |  |  |
| actin+Cytochala<br>sin D<sup>11</sup> | The reported site<br>corresponds to site<br>S4 | Ile136, Tyr169, Ala170,<br>Pro172, Met335, Phe375 | 2 - 20 μM | -8.33 -<br>-6.61 |
| actin+<br>Kabiramide C<sup>13</sup> | The reported site<br>corresponds to site<br>S4 | Ile-341, Ile-345, Ser-<br>348, and Leu-349, Tyr-<br>143, Gly-146, Thr-148,<br>Gly-168, Tyr-169, Ile-<br>345, Leu-346, Thr-351,<br>Met-355, and Phe-375 | 100 nM or<br>less | -9.60 |

## Conclusion

We present a detailed computational investigation of the binding behaviour of bacterial inhibitor A22 with eukaryotic actin protein using molecular docking, unbiased MD simulations, and infrequent well-tempered metadynamics simulations. From molecular docking and unbiased MD simulations, we showed that A22 predominantly binds in five binding sites in actin, including both previously reported sites (S1, S4) and new sites (S2, S3, S5).

From unbiased all-atom molecular dynamics simulations, we computed structural properties like RMSD, RMSF, R_g_ of the ligand-protein complexes at these five binding sites. Interaction analyses showed that A22 mostly forms transient interactions with actin protein except at sites S4 and S5, where long-lived interactions are present. In site S5, where a reduction in RMSF was observed in local stabilization at specific regions, the DNase-I binding loop region, other sites show structural stability. From iWT-MetaD simulations we characterize binding affinity, ligand dissociations, residence times, key gatekeeper residues at these specific-sites. we observed low binding affinities in S1-S4 sites (Δ*G* ∼ –2 to –3.4 kcal/mol), whereas in site S5 we observed highest binding affinity (Δ*G* ∼ – 6.05 ± 0.48 kcal/mol). The ligand residence times were in the range of ∼ µs to ms range, whereas for site S5 we observed residence time in order of ∼ 0.75 sec. Explicit analyses of the dissociation trajectories explore the dissociation pathways the ligand takes while dissociating from specific binding sites. Also, the key gatekeeper residues have been identified and reported. These provide deeper level mechanistic insights in the ligand dissociation processes and key protein residues involved in these processes. To see the relationship between the free energy and residence time, we correlated the free energy and residence time. It shows the free energy and residence time follows thermodynamics trend. However, we observed some deviations which might be due to enthalpy-entropy compensation effects. Our comparison with reported inhibitors in literature suggest A22 as a weak-action-binder with potential to modulate actin function through dynamics rather than tight-binding interactions.

Hence, this study provides a molecular-level insights into the actin-A22 interaction mechanism at its multiple binding sites, thermodynamics binding affinities, dissociation pathways, residence times and key gatekeeper residues. Therefore, this study provides a deeper understanding of cytoskeletal modulation by small molecules, computation framework for evaluating ligand unbinding mechanisms in multi-site protein targets. This molecular level detailed understanding might be advantageous to understand the effect of small molecule actin inhibitors on actin polymerization/depolymerization process and eventually leading to drug therapeutics.

## Data and Software Availability

The initial crystal structure of protein is available in the Protein Data Bank (PDB ID: 1YAG). The initial structure of A22 ligand is taken from Cc-MreB (PDB ID: 4CZG). The Gaussian 16 software was used for quantum optimization (https://gaussian.com/gaussian16/). Molecular docking was performed using AutoDock Vina 1.2.5 tool (https://vina.scripps.edu/) and VMD (https://www.ks.uiuc.edu/Research/vmd/), PyMOL (https://www.pymol.org/) were used for visualization purpose. All atom molecular dynamics simulations were carried out by using GROMACS-2022 software packages (https://manual.gromacs.org/2022/). Interaction fingerprint calculations were performed using ProLIF tool (https://github.com/chemosim-lab/ProLIF). Metadynamics simulations were performed using PLUMED (https://www.plumed.org/). Initial system coordinates, force field parameter files and input files for unbiased MD, infrequent metadynamics simulations are provided in the following link. (https://github.com/dpramanik-lab)

Various detailed information is available in the Supplementary Information.

## Supplementary Information

Table S1 provides actin binding sites, docked free energies, and amino-acid residues from docking simulations; Interaction fingerprints analysis criterion; Figure S1(a-e): interactions fingerprints of A22 for various binding sites; reaction coordinate and parameters for infrequent well-tempered metadynamics simulations; Figure S2: schematic representation of the selected groups of protein atoms and ligand atoms to construct RC; Figure S3: reaction coordinate (RC) over time plot and autocorrelation time of the RC; Figure S4: independent free energy profiles at different sites; Figure S5(a-e): time evolution of time derivative of acceleration factor; Figure S6(a-e): time evolution of RC (distance); Figure S7: KS test description and plots of KS test.

## Acknowledgements

A.K. thanks SRM University–AP for PhD fellowship. We thank the HPCC computing facilities provided by SRM University–AP supercomputer centre. D.P. thanks Science and Engineering Research Board (SERB), Government of India for supporting with the State University Research Excellence (SURE) project No. SUR/2022/004576. Author appreciates the infrastructure support received from SRM University AP (SRMAP/URG/GENERAL/2023-24/010), (SRMAP/URG/E&PP/2022-23/018), and (SRMAP/URG/GENERAL/2024-25/039).

We thank Sutharsan Govindarajan, Department of Biological Sciences, SRM University – AP for useful suggestions.

## Author Contributions

A.K.: computational work, analysing data, writing-review. D.P.: conceptualization, supervision, investigation, validation, visualization, writing-original draft, writing-review and editing. All authors read and approved the final manuscript.

